# Simulation of transcranial magnetic stimulation in head model with morphologically-realistic cortical neurons

**DOI:** 10.1101/506204

**Authors:** Aman S. Aberra, Boshuo Wang, Warren M. Grill, Angel V. Peterchev

## Abstract

Transcranial magnetic stimulation (TMS) enables non-invasive modulation of brain activity with both clinical and research applications, but fundamental questions remain about the neural types and elements it activates and how stimulation parameters affect the neural response. We integrated detailed neuronal models with TMS-induced electric fields in the human head to quantify the effects of TMS on cortical neurons. TMS activated with lowest intensity layer 5 pyramidal cells at their intracortical axonal terminations in the superficial gyral crown and lip regions. Layer 2/3 pyramidal cells and inhibitory basket cells may be activated too, whereas direct activation of layers 1 and 6 was unlikely. Neural activation was largely driven by the field magnitude, contrary to theories implicating the field component normal to the cortical surface. Varying the induced current’s direction caused a waveform-dependent shift in the activation site and provided a mechanistic explanation for experimentally observed differences in thresholds and latencies of muscle responses. This biophysically-based simulation provides a novel method to elucidate mechanisms and inform parameter selection of TMS and other forms of cortical stimulation.

## Introduction

Transcranial magnetic stimulation (TMS) is a technique for non-invasive modulation of brain activity using a magnetically-induced electric field (E-field)^1^. TMS is currently FDA-approved for cortical mapping and the treatment of several psychiatric and neurological disorders, and it is under investigation for many other indications^2^. However, rational design and optimization of TMS is impeded by our limited understanding of its neural effects. Fundamental questions persist regarding the neural types and elements that are activated, the spatial extent of activation, and how spatial and temporal parameters of TMS determine threshold and site of activation, particularly when considering the complexities of human brain geometry. Insight into the cortical origin and mechanisms of the physiological response to TMS has proven difficult: electrical recording techniques in the brain or spinal cord are invasive and mostly reserved for animal studies, generally suffer from long stimulus artifacts, as well as uncertainty about the stimulation focality, and have yet to produce a systematic assessment of the neural response across depths, positions, and cell types^3–5^.

Here, we model and quantify the responses of cortical neurons to TMS-induced E-fields and thereby address these questions about the interaction between TMS and neurons. Computational modeling is a powerful tool for investigating the mechanisms of TMS as well as for choosing stimulation parameters for more selective target engagement. Prior modeling efforts focused on calculating the spatial distribution of the induced E-field by TMS—typically using the finite element method (FEM) in head models derived from magnetic resonance imaging (MRI) data^6,7^. However, the spatial distribution of the E-field alone cannot predict the physiological effects of stimulation, and TMS can recruit distinct neural populations or elements based on different temporal dynamics of the E-field waveform (e.g. pulse shape, direction, width, and phase amplitude asymmetry)^8–12^. Therefore, the E-field must be coupled to neural models capturing the diversity and spatial distribution of the underlying neural elements, as well as their membrane dynamics, to quantify the neural response to stimulation.

To simulate the neural response to TMS, cable theory was adapted and implemented in increasingly detailed compartmental neuron models^13–18^. These studies concluded that axonal bends, branch points, and terminations are the most likely sites of action potential initiation by TMS. However, more recent studies suggest that TMS initiates action potentials at the cell body or axon initial segment^19–21^. Additionally, preliminary efforts to develop integrated models incorporating anatomically accurate E-field models with morphologically realistic neuron models lacked realistic axonal geometries or diversity of cell types and offered limited mechanistic insights^17,20^.

Our approach incorporated cortical geometry, neural membrane dynamics, and axonal morphology, all of which are critical to accurately model the effects of TMS. We adapted recently published cortical neuron models from the Blue Brain Project^22^ to the geometry of mature human cortical neurons, including excitatory and inhibitory cell types across all cortical layers, realistic axon morphologies, experimentally validated electrophysiological properties, and morphological variability within cell types^23^. We embedded these model neurons in cortical layers within the gray matter of a realistic FEM model of a human head. This level of detail and diversity of model neurons achieved a more biologically-plausible model of the direct cortical response to TMS than any previously published. Finally, we used this integrated model to simulate the neural response to TMS of primary motor cortex (M1) with several E-field directions and pulse shapes, including conventional and controllable pulse parameter TMS (cTMS). The simulations answered three key questions that have been controversial in the field: 1) which neural types and elements does TMS activate, 2) what is the site of activation by TMS of the motor cortex, and 3) how is the site of activation affected by E-field direction and waveform?

## Results

### Threshold and directional sensitivity vary by cell type

We quantified variations in excitation threshold and intrinsic sensitivity to local E-field direction for each model neuron, independent of their cortical location and the E-field non-uniformity in the head model (Figure 1A–D). For all E-field directions, the site of maximal depolarization and action potential initiation was an axonal termination aligned with the E-field. Consequently, the densely-branching axonal arbors exhibited numerous potential activation sites, and the orientations where specific axonal branches aligned with the E-field in each model neuron corresponded to local minima in the threshold–direction maps.

**Figure 1.**
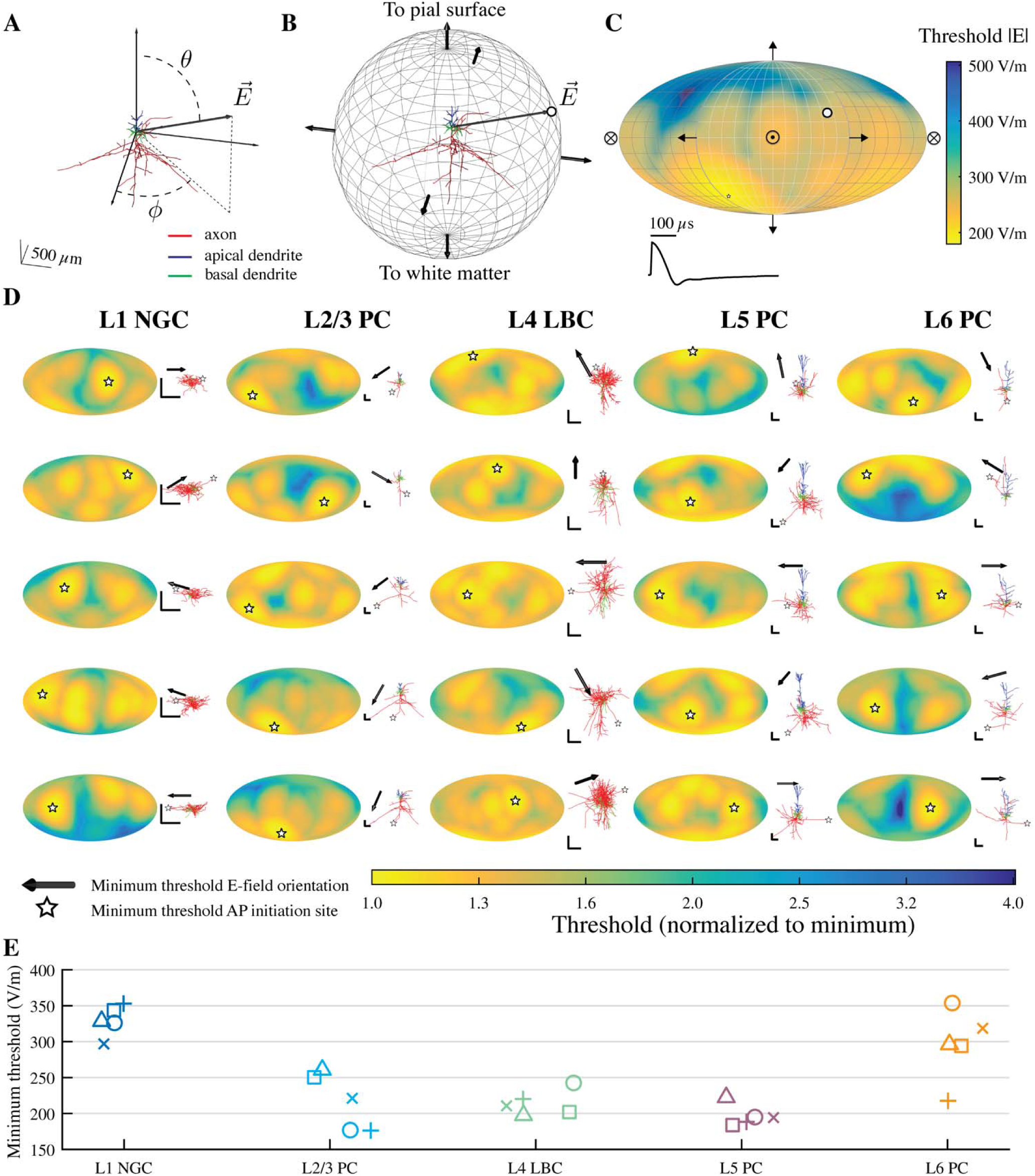
Threshold dependence on cell type and direction of uniform E-field. **A)** Coordinate system shown for example cell (L2/3 PC). Somato-dendritic axis aligned to z-axis, with polar angle *θ* and azimuthal angle *Φ*. **B)** Uniform E-field directions represented as normal vectors on sphere centered at origin. Thresholds were calculated for 398 directions spanning the sphere, and each threshold value was represented as a point on the sphere (white dot) corresponding to the E-field vector 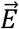 **C)** 3D threshold–direction map projected into 2D using Mollweide projection. White dot indicates threshold value for example vector 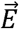 in **B**, crossed circle represents E-field pointing into the page, and circle with dot represents E-field pointing out of the page. Bottom: Recorded MagPro X100 Monophasic TMS waveform. **D)** Threshold–direction maps for all cell types and their virtual clones, i.e. models of the same cell type with stochastically varied morphologies, normalized to the minimum for each clone. Within cell type, threshold–direction maps for each clone are ordered by minimum threshold, with lowest minimum threshold at the bottom. White star denotes minimum threshold orientation. Corresponding cell morphology plotted to the right of each map with same color scheme as in **A**. Black arrow points in direction of minimum threshold orientation, matching white star. All scale bars are 250 µm. **E)** Box-plots of minimum threshold for 5 clones of each cell type, listed by layer.

Thresholds varied substantially both within and between cell types. Within cell type, the minimum threshold amplitudes varied by 18%–73% relative to the lowest threshold clone (Figure 1E). Based on the median within each cell type, the minimum threshold amplitudes were lowest for the layer 5 (L5) pyramidal cell (PC), followed by L4 large basket cell (LBC), L2/3 PC, L6 PC, and L1 neurogliaform cell (NGC) (Figure 1E). Notably, the minimum threshold among the L2/3 PCs was lower than that of the L5 PCs, despite the higher median.

Threshold anisotropies were used to characterize directional sensitivity and were higher for the pyramidal cell types (2.29–5.09) compared to the interneurons (1.73–3.45) (Figure 1D). The higher threshold anisotropies in the pyramidal cells reflected their more asymmetric, elongated axons relative to the more spherically-symmetric axons of the interneurons, which have a broader distribution of axon branch orientations. The mean threshold differences between transverse and inward or outward E-fields, relative to the somatodendritic axis of each model neuron, ranged from −34% to 32% (positive value indicates higher threshold for transverse E-field) (Supplementary Figure 1). L5 PCs had the strongest preference for both inward and outward E-field directions, relative to transverse, followed by the L2/3 PCs and L4 LBCs, while the L1 NGCs and L6 PCs generally had lower thresholds for transverse E-field directions (Figure 1D; Supplementary Figure 1).

In addition to these well-characterized and validated models^23^, we also applied the same age- and species-related modifications to the entirety of the model neurons in the Blue Brain library and determined their minimum thresholds for activation with uniform E-fields. The inhibitory basket cell types in L2/3, L5, and L6 had thresholds that were comparable to the L4 LBCs; similarly, the subset of pyramidal cell types we selected were representative of the thresholds of the remaining pyramidal cell types within their respective layers (Supplementary Figure 2). Thus, we proceeded with the originally selected set of model neurons for quantifying the effects of TMS.

### TMS activates cortical layers 2–5 in gyral crown

We simulated patterns of neural activation by populating the head model with model neurons in a region of interest (ROI) comprising the left motor hand knob in M1 and positioning the TMS coil in an orientation corresponding to the lowest threshold for evoking motor activity (Figure 2). The E-field distribution in the brain was approximately tangential to the scalp, with the largest amplitudes at gyral surfaces perpendicular to the E-field direction, and decayed rapidly with depth into the sulcus (Figure 3A,B).

**Figure 2.**
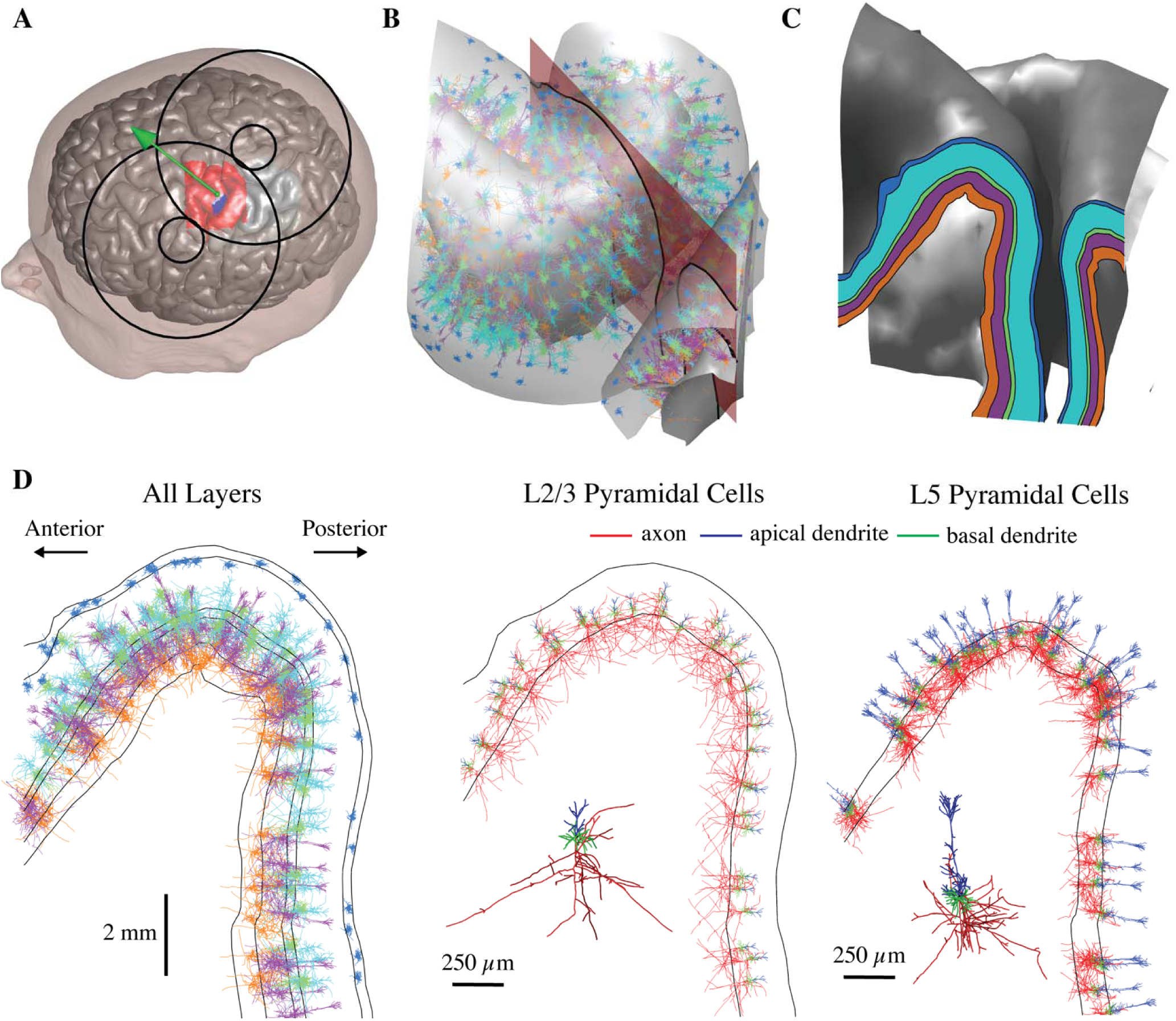
Embedding populations of cortical neuron models in FEM models of TMS induced E-fields. **A)** Scalp and gray matter meshes are shown with the overlying TMS coil outline The coil center and orientation are given by the green sphere and arrow, the hand knob region populated with neurons is indicated in red, and the putative FDI representation used in Figure 6 is shown in blue. **B)** Model neurons located in the crown of the pre-central gyrus between the gray matter and white matter surfaces. One clone from each layer is shown with color corresponding to layer (shown in **C**); five co-located model neuron populations (virtual clones of each cell type) are simulated in each layer. The 2D analysis plane (red) is used to visualize threshold data in Figure 5 and also shown in **C**–**D**. The plane is parallel to coil orientation (45° relative to midline). **C)** Cortical layers used to place and orient model neurons shown in 2D analysis plane, extracted by intersecting the analysis plane with the layer surfaces. **D)** Neural populations from **B** visualized within the 2D analysis plane. Neurons in all five layers are plotted with their respective layer colors (left). L2/3 (middle) and L5 (right) PC populations are plotted with axon, apical dendrites, and basal dendrites colored separately.

**Figure 3.**
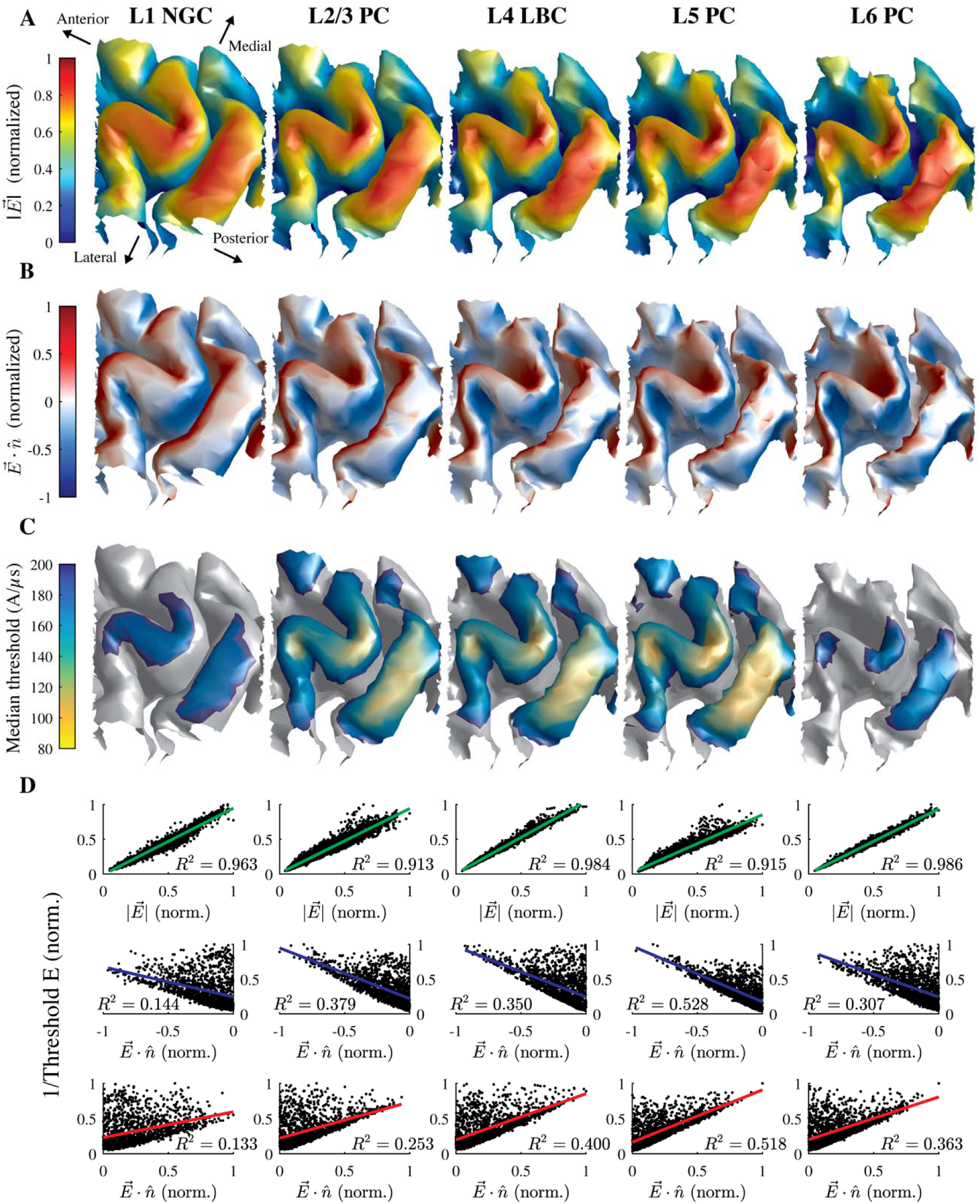
Layer-specific spatial distribution of activation thresholds correlate better with the E-field magnitude than normal component. **A)** Magnitude of simulated E-field (normalized across layers) on layer surfaces for L1– L6 arranged adjacent to each other. **B)** Component of E-field normal to layer surfaces (normalized across layers). Positive values indicate E-field pointing out of surface and negative values indicate inward E-field. **C)** Median thresholds (across 5 clones and 6 rotations) for monophasic P–A simulation. **D)** Inverse threshold of each cell plotted against E-field magnitude (top) and absolute value of normal component (bottom) at soma, both normalized to maximum within layer (in **A**). Each plot includes *R*^2^ value for linear regression of inverse threshold with corresponding E-field metric.

Monophasic stimulation with dominant E-field induced in the posterior–anterior (P–A) direction generated the lowest thresholds in L5 neurons, followed by L2/3 and L4, L6, and L1 (Figure 3C, also Figure 6). The recruitment order matched that of uniform E-field stimulation, with some degree of overlap between L2/3 and L4, and activation also occurred exclusively at the terminals of the axon collaterals (Figure 4).

**Figure 4.**
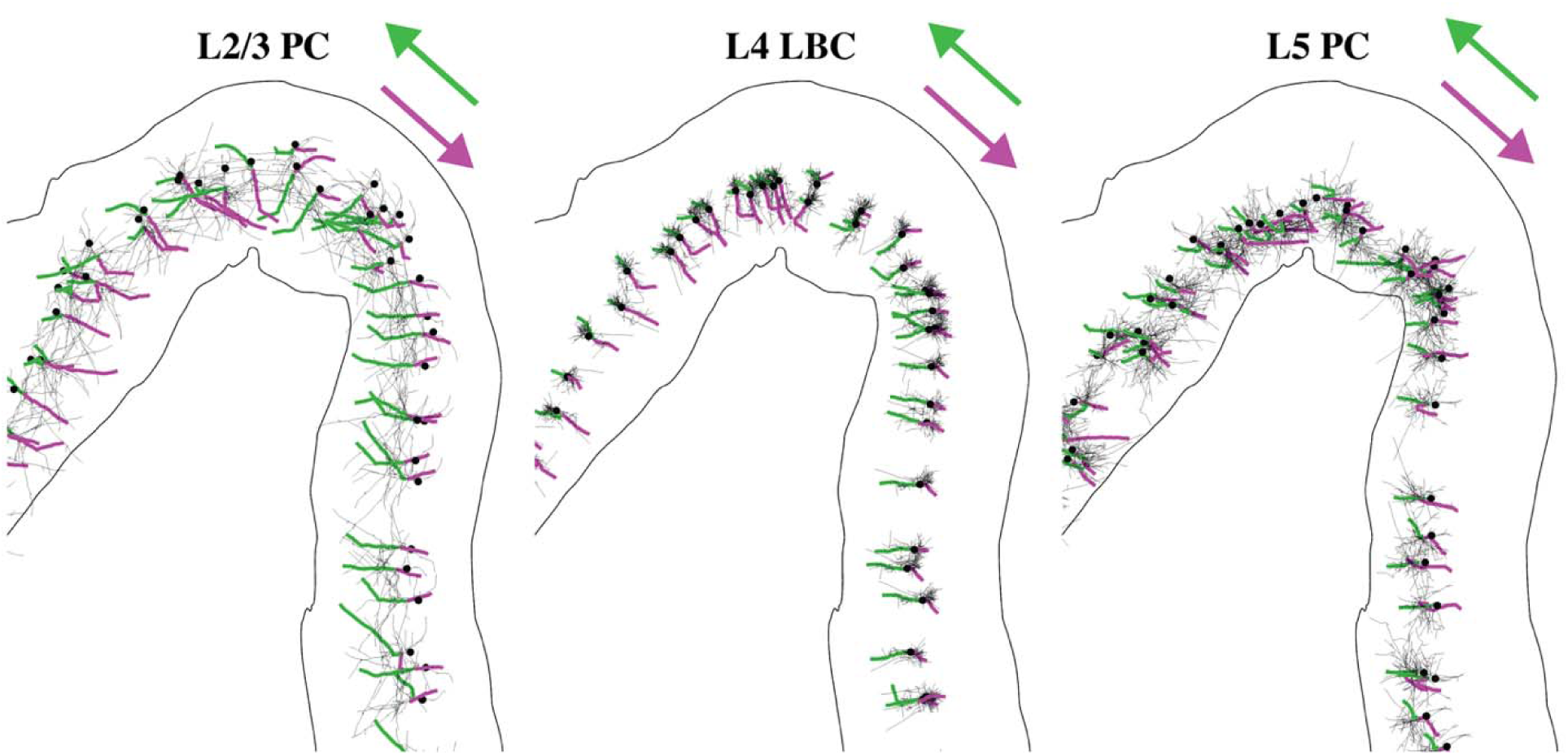
TMS activates axonal terminations aligned to local E-field direction. Axonal arbors for single population of L2/3 PCs, L4 LBCs, and L5 PCs with directly activated branch colored from AP initiation point (terminal) to proximal branch point for monophasic P–A (green) and A–P (magenta) stimulation. Dendrites not shown. Somas indicated by black dots

**Figure 5.**
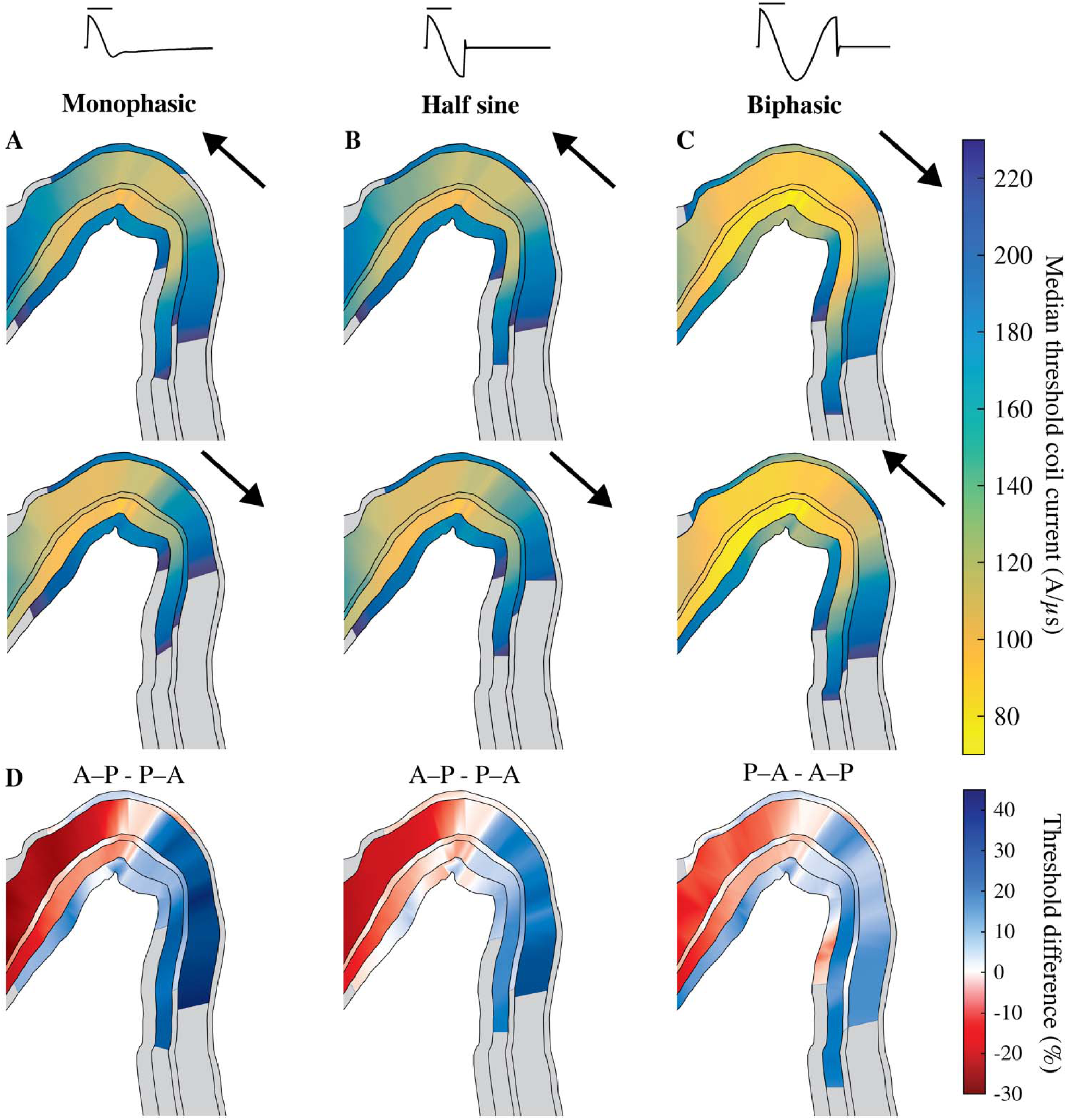
Layer-specific spatial distribution of activation varies with TMS pulse waveform and direction, shown in a cross-section of the hand knob (as in Figure 2C). Median thresholds for L1–L6 on analysis plane through pre-central gyrus, parallel to coil handle and near coil center for **A)** monophasic, **B)** half sine, and **C)** biphasic stimulation in the P–A and A–P directions. Arrows indicate direction of initial phase of E-field waveform. Note that biphasic stimulation conditions are plotted in opposite order to group stimuli by the direction of their dominant waveform phase. **D**) Percent difference in median thresholds between P–A and A–P current directions, indicated in title. Regions where thresholds for both P–A and A–P were above 230 A/µs (maximum in **A–C**) are colored gray.

**Figure 6.**
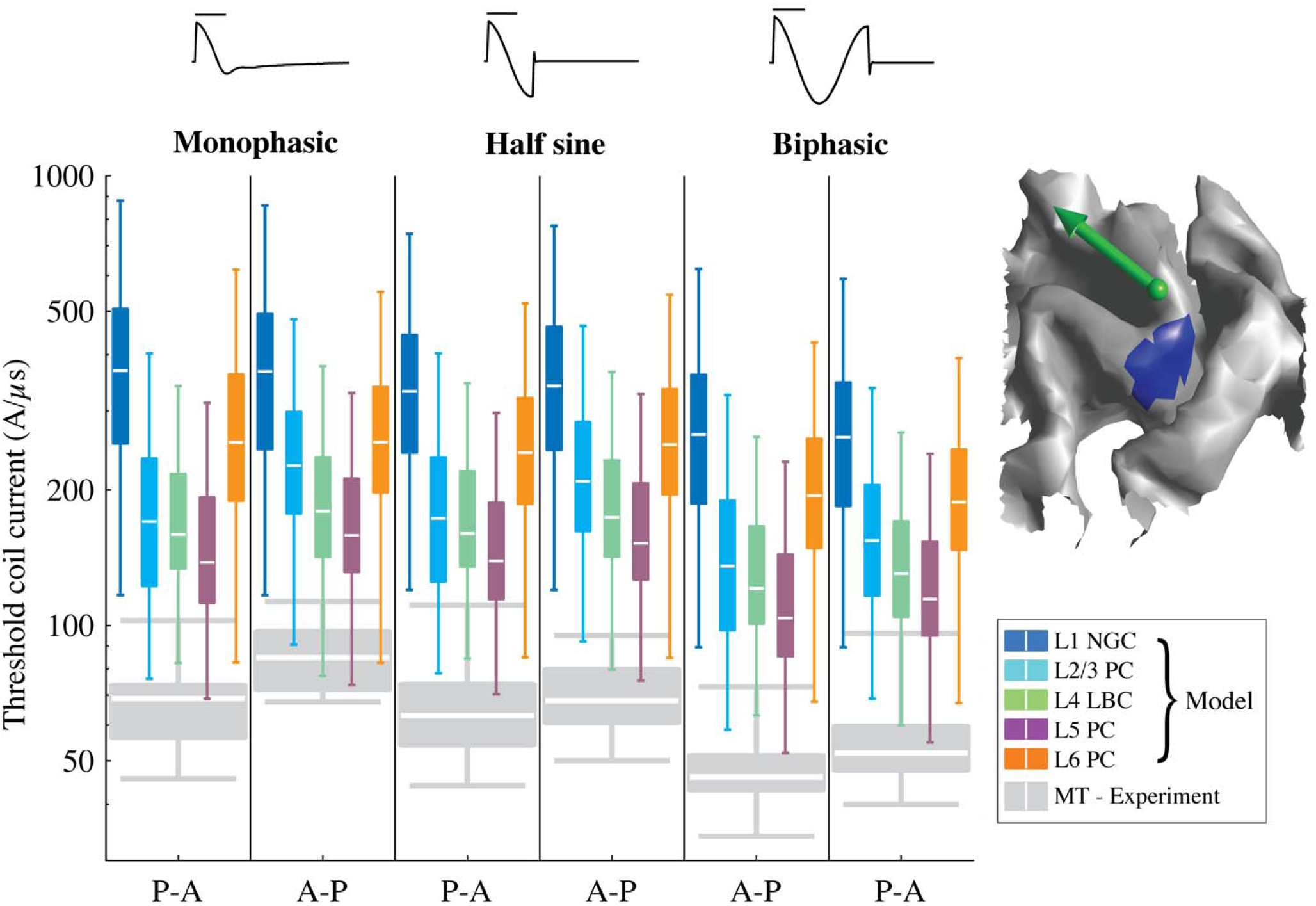
Activation thresholds within approximate FDI representation in hand knob. Box-plots of model thresholds for each pulse waveform and direction combination are shown in log scale. FDI on L5 surface is marked in blue (top right). Each box-plot includes thresholds from 5 clones and 6 rotations at each position in the FDI representation. Green arrow indicates center of TMS coil and direction of induced current (for P–A stimulation). Box-plots of experimental motor threshold (MT) data26 (12 subjects) included in gray.

Previous publications hypothesized that the threshold for neural activation by TMS is inversely proportional to either the E-field magnitude^7^ or its normal component relative to the cortical surface, based on the orientation of cortical columns^24,25^, thereby generating the lowest thresholds in either the gyral crown or the sulcal wall, respectively. Thresholds were lowest in the pre- and postcentral gyral crowns for all five layers, corresponding to regions of maximal E-field amplitude. The low threshold regions also extended to the gyral fold lateral to the hand knob, where the E-field magnitude remained high relative to the peak under the coil center, as well as deeper into the posterior wall of the central sulcus relative to the anterior wall (Supplementary Figure 3). Thresholds within the sulcal walls were substantially higher than the more superficial gyral regions, and activation reached relatively deeper into the sulcal wall for the low-threshold middle layers: L2–L5.

Throughout the ROI, the thresholds of all cell types were more strongly correlated with the local E-field magnitude (*R*^2^ ≥ 0.913) than the normal component (inward: *R*^2^ ≤ 0.528; outward: *R*^2^ ≤ 0.518) (Figure 3D), indicating that the E-field magnitude was a markedly stronger driver of neuronal activation than the normal component.

### Activation depends on stimulus waveform and direction

Shifts in action potential initiation sites with current direction led to layer- and waveform-specific shifts in the spatial distributions of thresholds. For monophasic P–A stimulation, the activated terminals in the low-threshold L2–L5 were located in the downward projecting axonal branches in the gyral lip and sulcal wall and the horizontal and oblique collaterals in the gyral crown (Figure 4). For reversed monophasic stimulation with anterior–posterior (A–P) direction, activation shifted to an axonal terminal in the opposite direction for each model neuron (Figure 4).

Using the same current direction, monophasic, half sine, and biphasic pulses produced similar threshold distributions, with the lowest thresholds in the gyral crown and lip (Figure 5A–C). Switching from P–A to A–P monophasic TMS resulted in an anterior shift of activation in L2–L5 of the pre-central gyrus; in the analysis plane, thresholds were up to 30% higher on the posterior lip and up to 45% lower on the anterior lip (Figure 5D). In contrast, for the half sine and biphasic waveforms, which have similar peak E-field strengths in each phase, the shifts in activation were in the same direction, but substantially reduced. Note that for biphasic pulses, the second phase with longer duration and direction opposite to the initial phase is the dominant one, and therefore the effect of directionality is reversed^16,26^.

Linear regressions between the thresholds of the model neurons within the putative first dorsal interosseous (FDI) representation^7^ and experimentally measured motor thresholds (MTs)^26^ (Figure 6) across all waveforms and directions yielded strong correlations in all layers (*R*^2^ > 0.75; *p* < 0.03), with the strongest correlations in L2–L5 (*R*^2^ > 0.85) (Supplementary Figure 4). When the current direction was reversed from P–A to A–P, the L1 and L6 neurons exhibited less than a 5% change in threshold for all pulse waveforms. In contrast, for the L2–L5 neurons, the threshold increased by 8–33% for all pulse waveforms, with a greater increase for the monophasic pulse than for the half-sine and biphasic pulses, consistent with the experimental data^26^ (Figure 6).

### Model strength–duration time constants match experiments

Time constants of single model neurons exhibited slight variations throughout the ROI for the L4 LBCs and L5 PCs, while the L2/3 PCs had consistently longer time constants in the posterior-facing sulcal walls throughout the ROI (Figure 7A). Using median thresholds in the FDI representation, the strength–duration curves for L2–L5 neurons agreed remarkably well with experimental measurements (Figure 7B). The proportion of activated cortical neurons corresponding to MT is unknown, but the slight variation of time constant showed insensitivity to the proportion of activation and overall agreement with the experimental estimate (200 ± 33 µs; mean ± SD)^11^ (Figure 7C). L2/3 PCs had longer time constants (224 – 288 µs) than model neurons in the other layers and the experimental estimate, and this was consistent with the markedly increased time constants in the sulcal wall as estimated using single neuron thresholds (Figure 7A).

**Figure 7.**
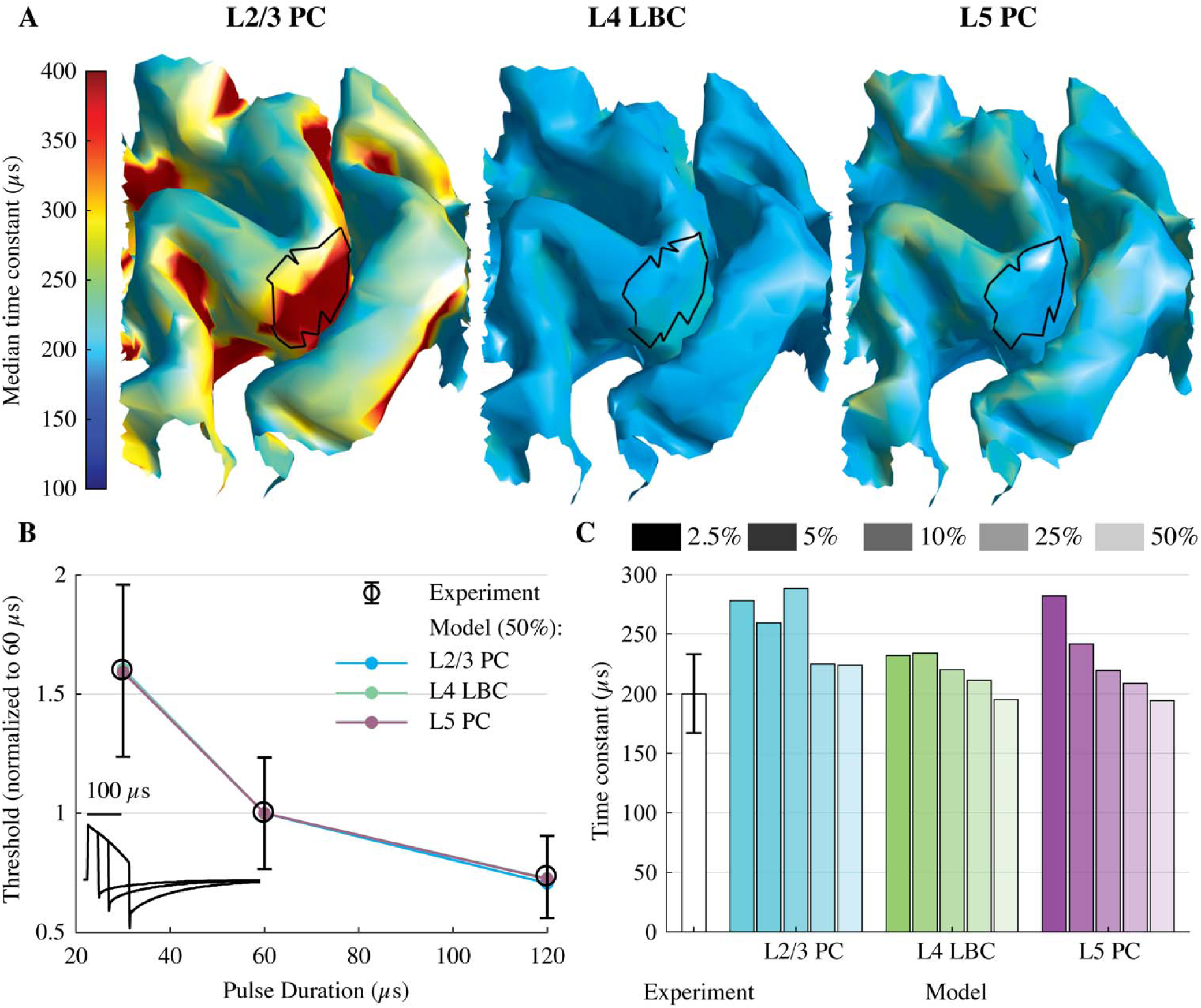
Strength–duration time constants of model neurons match experimental measures. **A)** Median time constant (across 5 clones and 6 rotations) for layers with lowest activation thresholds, estimated for each model neuron using their activation thresholds for 30, 60, and 120 µs cTMS pulses. Assumed FDI representations are outlined in black. **B)** Strength–duration curves using median neuronal population threshold within the model FDI representation and experimentally measured mean (± SD) motor thresholds^11^. cTMS waveforms are shown in the bottom left corner. **C)** Experimentally estimated time constant^11^ (mean ± SD) and model time constants for a range of cutoff threshold percentiles (2.5–50%) within the FDI representation.

### Discussion

We developed a detailed, multi-scale model of the cortical response to TMS by populating an MRI-derived FEM head model with realistic models of neurons across the cortical layers. The neuron models approximated the geometry of human cortical cells and included extensive axonal arbors, which were necessary to predict the dependence of activation on stimulation direction in each cell type. The results revealed several important conclusions regarding the neural mechanisms of TMS of the motor cortex. TMS exclusively activated neurons at their axon terminals aligned to the local E-field direction. Due to their complex axonal arborizations, neurons were activated by E-fields both normal and tangential to the cortical surface, and the E-field magnitude was a much stronger predictor of neural activation than the E-field component normal to the cortical surface. In both the uniform E-field threshold–direction maps and the multi-scale model, thresholds varied substantially across neurons in different cortical layers, in a manner that did not simply reflect their respective distances from the coil: the L5 PCs had the lowest thresholds, followed by L2/3 PCs, L4 LBCs, L6 PCs, and L1 NGCs. TMS of M1 activated with lowest intensity neurons in the crown and lip of the precentral gyrus due to the higher E-field magnitude there as compared to the sulcal wall. Reversing the current direction from P–A to A–P caused an anterior shift in the threshold distributions of L2/3–L5, with the strongest effects in the pyramidal cell types and for the monophasic waveform. These trends agreed remarkably well with variations in MT observed in human subjects.

TMS preferentially activated axon terminals in all cell types, in both the uniform and non-uniform E-fields. As such, our results differ from two widely cited modeling studies which concluded that TMS initiates action potentials at the soma or initial segment^19,20^. Indeed, their results can be explained by implementation errors in the coupling between the E-field and the neural cable models^15^. Activation at the soma or dendrites is unlikely due to their long membrane time constants (> 5 ms) compared to myelinated axons (< 400 µs)^27,28^. These axonal time constants agree with strength–duration time constants measured for TMS of the motor cortex^11,29^, and were reproduced by our models. In contrast, direct polarization of cell bodies, before an axonal compartment was activated by suprathreshold TMS pulses, was well below action potential threshold (≤ 2–3 mV). Further evidence for preferential activation of axonal terminals comes from experimental data on the corticospinal I_1_-wave^30^, which is thought to originate from excitatory, monosynaptic inputs to the pyramidal tract neurons projecting directly to hand motoneurons^31^. The I_1_-wave threshold is relatively unaffected by voluntary contraction, GABAergic drugs, or the paired-pulse paradigm known as short-interval intracortical inhibition^30^. This is expected if the pre-synaptic inputs to PTNs were activated at their distal axon terminals, where the membrane potential is less affected by synaptic inputs than the dendrites, soma, or axon initial segment. Thus, our finding that TMS preferentially activated axon terminals, while inconsistent with some prior modeling studies, is consistent with complementary experimental results.

Including realistic axonal geometries is critical to predicting accurately the variations in threshold and directional sensitivity between and within cell types, and this had significant implications for the mechanisms of neural activation in the full model. Previous biophysical models with idealized, straight axons found extremely high threshold anisotropies and could not predict variations in collateral activation between cell types^16,20^. On the other hand, we observed significant variations in activation thresholds both between and within cell types as a result of axonal geometry. In addition, the phenomenological cortical column cosine model argues that pyramidal cells are depolarized by E-field directed into the cortical surface (parallel to the cortical columns), while interneurons are depolarized by tangential fields^24,25,32^. This led to the prediction that TMS activates pyramidal cells in the sulcal wall, where the E-field is directed into the cortical surface, and therefore, neural activation should be proportional to the normal component of the E-field^24,25,32^. While the low-threshold cells in our model (L2–5) exhibited a preference for normal relative to transverse E-field, this effect was relatively small, with typical threshold differences of less than 30% (Supplementary Figure 1). Since the E-field activated the terminals of aligned axonal branches, but not cell bodies or dendrites, the dense local and distal axon collateralization of cortical PCs and interneurons enabled activation for all E-field directions^23^. Consequently, the spatial distribution of thresholds for a given coil configuration was much more highly correlated with the E-field magnitude than its normal component. Neural activation is therefore more likely to occur in the gyral crown and lip, which are exposed to larger E-field magnitudes, than the sulcal wall. These results provide a clear mechanistic explanation of recent studies relating MTs for a range of TMS coil orientations and positions with E-field distributions calculated in subject-specific head models^6,7^.

The integrated model enabled analysis of waveform- and direction-dependent effects that would be indiscernible using the E-field distribution alone. TMS of the motor cortex produces motor evoked potentials (MEPs) with 2–3 ms longer latencies for current in the A–P than in the P–A direction^8,26,29,33,34^. In the model, reversing the current direction for the monophasic waveform from P–A to A–P produced an anterior shift in the spatial distribution of activation of L2/3 and L5 PCs. These shifts with current direction were explained by the uniform E-field threshold–direction maps for these cell types, as they indicated slight preferences for E-fields oriented into the cortical surface for L2–5 neurons (Supplementary Figure 1). While this preference was not sufficient to reduce thresholds in the sulcal wall, where the E-field is nearly perpendicular to the cortical surface, it did alter thresholds within the gyral crown and lip dependent on the current direction. This waveform-dependent shift in the region of cortical activation could explain the longer MEP latencies of monophasic A–P stimulation relative to P–A stimulation, which is not observed for the more symmetric waveforms. Corticomotoneuronal cells that make monosynaptic connections with alpha motoneurons are found mostly in the anterior bank of the central sulcus^35^. Based on our model, P–A stimulation would therefore mostly activate the corticomotoneuronal cells monosynaptically, producing MEPs with shorter latencies, while A–P stimulation would activate rostral M1 or pre-motor pyramidal cells producing MEPs polysynaptically with longer latencies. In fact, excitatory inputs to M1 from ventral and dorsal pre-motor cortex were identified in monkeys^36–38^ and humans^39^ with conduction latencies matching late corticospinal volleys generated by single-pulse TMS (I-waves)^40^, suggesting they may be recruited by A–P stimulation^30^. These results support that A–P and P–A stimulation of M1 activate different sets of cortical neurons that indirectly generate downstream corticospinal activity, rather than different sites within the same neurons, as argued elsewhere^30^.

Although a substantial advance over prior work, several limitations of the model should be noted. Obtaining realistic threshold amplitudes in neural models of TMS is a persistent challenge for the field^16–18^. Sommer et al. reported MTs for monophasic P–A stimulation between 45.6–102.6 A/µs for subjects at rest. These values are close to the minimum threshold of L5 PCs in the FDI representation within our model (68.7 A/µs); however, sub-MT stimuli are known to activate intracortical circuits^41^, suggesting that model thresholds may still be too high. Uniform overestimation of the activation thresholds may be related to limitations in the axon models, specifically in the sodium channel model, as well as differences in morphological features related to age, species, and brain region^23^.

Another limitation was the lack of corticospinal models extending into the white matter, which may be activated directly by TMS with certain coil configurations^30^, as well as medium to long-range intracortical axonal projections^42^. The trajectories of corticofugal projections can be incorporated using diffusion tensor imaging (DTI) tractography^43,44^, whereas obtaining intracortical projections is more challenging. We focused on modeling a single excitatory or inhibitory cell type per layer, but there were additional cortical cell types with low thresholds for activation that we did not simulate in the full multi-scale model. Our supplementary simulations suggest inhibitory basket cells in L2/3, L5, and L6 may be recruited by TMS at similar amplitudes to the L4 large basket cells. We also focused on direct activation by TMS and did not include synaptic connections between neurons. Network models with both reconstructed morphologies and realistic simulations of the E-field pose a daunting computational task, but may be important in modeling short-term circuit dynamics in response to TMS.

Finally, methods for computing the TMS-induced E-field in FEM models have limitations^45^, including the assumption of homogeneous, isotropic conductivities in gray matter and the sharp conductivity boundary at its border with the white matter^45^. In our model, the unnatural E-field gradients did not produce activation in L5/6 PC axons that crossed the gray–white matter boundary, suggesting the latter issue is not critical. Additionally, we used a single subject’s head model, and sites of activation may vary with cortical geometry^46^.

### Outlook

This work introduces a computational modeling platform for simulating the response of cortical neurons to transcranial magnetic stimulation and enables quantification of the effects of pulse waveforms, coil configurations, and individual head anatomies on the cell-type specific cortical response. Our results demonstrate the importance of incorporating non-linear neural membrane dynamics and realistic axon morphologies to capture the effect of the stimulation parameters. This platform will allow continued mechanistic studies and optimization of TMS and other cortical stimulation technologies to guide the rational design of non-invasive brain therapies.

## Methods

### Neuron models

The neuron models were modified versions of the multi-compartmental, conductance-based models implemented by the Blue Brain Project^22,47^ in the NEURON v7.4 simulation software^48^. The original models include 3D, reconstructed dendritic and axonal morphologies of cell types from all 6 cortical layers, with up to 13 different published Hodgkin-Huxley-like ion channel models in the soma, dendrites, and axon initial segment. Each cell type had five clones, which were generated by introducing stochastic variations in their dendritic and axonal geometries to reflect morphological diversity within cell type. Previously, we adapted a subset of these model neurons to the biophysical and geometric properties of adult, human cortical neurons and characterized their response to stimulation with exogenous E-fields^23^. These modifications included scaling the morphologies to account for age and species differences, as well as myelinating the axonal arbors, scaling ion channel kinetics to 37° C using a *Q*_10_ of 2.3, and assigning ion channel properties to the entire axon arbor. The final set of cell types included the layer 1 (L1) neurogliaform cell with a dense axonal arbor (NGC), L2/3 pyramidal cell (L2/3 PC), L4 large basket cell (L4 LBC), L5 thick-tufted pyramidal cell with an early bifurcating apical tuft (L5 PC), and L6 tufted pyramidal cell with its dendritic tuft terminating in L4 (L6 PC). The pyramidal cell types were selected based on their hypothesized involvement in I-wave generation^31^, and the L1 NGC cell type was selected based on its hypothesized involvement in TMS-induced inhibition^49^. Since myelin is expected to reduce thresholds^23^, the LBC cell type was selected based on the finding that the vast majority of myelinated, inhibitory axons belong to basket cells^50^. Further details can be found in our previous publication^23^ and these models can be downloaded from ModelDB (https://senselab.med.yale.edu/modeldb/ShowModel.cshtml?model=241165).

We made an additional modification to the L5 and L6 PC models to account for the truncation of their main axons in the slicing process. The main axons of L5 and L6 PCs project subcortically in mature animals, while their collaterals can extend both locally and for several millimeters horizontally^51^. The truncated main axon terminals in these model neurons were therefore unrealistically close to their cell bodies. To exclude them from activation, we disabled these main axon terminals by setting the terminal compartment diameter to 1000 µm. We quantified the effect of disabling the main axon terminals of the L5 and L6 PCs: leaving the main axons intact decreased the thresholds for downward E-fields for at least three of the five L5 PCs, reduced activation of horizontal collaterals and increased threshold anisotropy (Supplementary Figure 5). Similar effects were observed with the L6 PCs (Supplementary Figure 6).

### Head model for electric field simulation

Induced E-fields were computed in the realistic volume conductor head model (Figure 2A) provided as default in SimNIBS v2.0, an open-source simulation package that integrates segmentation of magnetic resonance imaging (MRI) scans, mesh generation, and finite element method (FEM) E-field computations^45^. The FEM model was generated using the T1- and T2-weighted images and segmentation from the SimNIBS example data set of a healthy subject, which included white matter, gray matter, cerebrospinal fluid, bone, and scalp tissue volumes. The MR images were acquired from a healthy subject with the approval of the Ethics Committee of the Medical Faculty of the University of Tübingen^52^. The white matter layer was assigned anisotropic conductivity using the diffusion tensor imaging (DTI) data and the volume normalized approach, with the mean conductivity of each tensor scaled to match the isotropic conductivity of 0.126 S/m^43^. The remaining four tissue volumes were assigned isotropic conductivities as follows (in S/m): gray matter: 0.276, cerebrospinal fluid: 1.790, bone: 0.010, scalp: 0.250 (according to Bungert et al.^7^). The final mesh comprised approximately 200,000 nodes and 3.6 million tetrahedral elements. Further details on the FEM model generation process were described by Windhoff et al.^52^. E-field distributions were computed with the SimNIBS models of the MagVenture MC-B70 figure-of-8 coil (P/N 9016E056, MagVenture A/S, Denmark) or the Magstim 70 mm figure-of-8 coil (P/N 9925-00, Magstim Co., Spring Gardens, Whitland, Carmarthenshire, UK), for a coil-to-scalp distance of 2 mm and a coil current of 1 A/µs. The MagVenture coil was used for simulations with conventional TMS waveforms to match the experimental setup of Sommer et al.^26^, and the Magstim coil was used for simulations with cTMS1 waveforms to match the setup of Peterchev et al.^11^. We simulated TMS of the left hand motor area by positioning the coil over the motor hand knob, located on the precentral gyrus^53^. Following the convention for TMS of M1, the coil was oriented to induce currents perpendicular to the central sulcus, directed 45° relative to the midline (Figure 2A); for monophasic P–A stimulation, this orientation corresponds to the lowest threshold for evoking motor activity, measured by motor evoked potentials (MEPs) in the first dorsal interosseous (FDI) or abductor digiti minimi, although there is considerable inter-individual variability^54,55^.

### Embedding neurons in head model

To generate layer-specific populations of neurons, surface meshes representing the cortical layers were interpolated between the gray and white matter surface meshes at normalized depths (the total depth of gray matter is 1): L1: 0.06, L2/3: 0.4, L4: 0.55, L5: 0.65, L6: 0.85 (Figure 2C). These depths were estimated from primate motor cortex slices, in which the boundaries between adjacent layers were at normalized depths of 0.08 (L1–L2/3), 0.51 (L2/3–L4), 0.59 (L4–L5), and 0.81(L5–L6)^56^. M1 is traditionally thought to lack a layer 4^57^, but it was included here based on studies in both mice and primates that demonstrated a functional layer 4 in motor cortex with canonical inputs from thalamus and intra-columnar outputs to L2/3^58–60^.

A 32×34×50 mm^3^ region of interest (ROI) containing the M1 hand knob on the pre-central gyrus and the portion of the postcentral gyrus opposite to it was selected for populating with model neurons (shown in red in Figure 2A). Layer surface meshes were discretized with 3000 triangular elements per surface, yielding a density of approximately 1.7 elements per mm^2^. Single model neurons were placed in each of the 3000 elements by centering the cell bodies within the elements (Figure 2B). Based on typical columnar structure^61,62^, the model neurons were oriented to align their somatodendritic axes with the element normals and randomly rotated about their somatodendritic axes in the azimuthal direction (see Figure 1A for definition of spherical coordinates). Each model neuron was simulated for five additional azimuthal rotations in 60° increments to sample the full range of possible orientations, and the five clones of each cell type were co-located within each element, increasing the effective cell density thirty-fold to approximately 50.7 cells/mm^2^ in each layer. Figure 2D depicts, in a cross-section of the hand knob, the placement of a single clone of each cell type in all five layers as well as the L2/3 and L5 PC populations plotted alone with different colours assigned to axons, apical dendrites, and basal dendrites. Mesh generation, placement of neuronal morphologies, extraction of E-field vectors from the SimNIBS output, NEURON simulation control, analysis, and visualization were conducted in MATLAB (R2017a, The Mathworks, Inc., Natick, MA, USA).

### Coupling electric fields to neuronal simulations

Using the quasi-static approximation^63,64^, the TMS induced E-field was separated into its spatial and temporal components. The cable equation has been adapted for magnetically induced E-fields to one-dimensional compartmental models^14,15,65^. The spatial component of the exogenous E-fields can be applied to cable models in NEURON using the `extracellular` mechanism^48^, which requires expressing the E-field in terms of an extracellular scalar potential at each compartment. However, magnetic induction produces a non-conservative E-field distribution that cannot be converted to scalar potentials. To couple the E-fields computed in the FEM simulations of TMS to the cable models, *quasipotentials*^15,17^ were calculated for each cell by numerically integrating the E-field component along each neural process

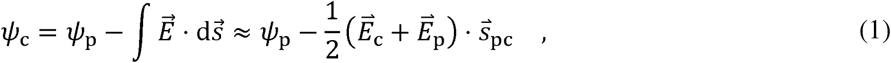

where *Ψ* is the extracellular quasipotential, 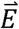 is the electric field vector, 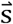 is the displacement vector, and subscripts c and p indicate the current/child compartment and previous/parent compartment in the tree structure with the soma arbitrarily designated as the root and reference point (*Ψ*_soma =0)_. The quasipotentials were calculated at the compartment centers, and then directly substituted as extracellular potentials in NEURON.

With the spatial distribution of quasipotentials determined, the temporal component of the E-field was included by uniformly scaling the distribution over time by various current waveforms. Monophasic, half sine, and biphasic waveforms generated by a MagPro X100 stimulator (MagVenture A/S, Denmark) with a MagVenture MCF-B70 figure-of-8 coil (P/N 9016E0564) were recorded using a search coil and sampling rate of 5 MHz (shown in Figure 5 and Figure 6). We also used waveforms generated by a cTMS1 device with a Magstim 70 mm figure-of-8 coil for 30, 60, and 120 µs pulse widths (shown in Figure 7B)^11,66^. The TMS E-field waveforms were down-sampled with 5 μs time steps for computational efficiency, and normalized to unity amplitude for subsequent scaling in the neural simulations.

As in our previous study^23^, the neural models were discretized with isopotential compartments no longer than 20 µm and solved using the backward Euler technique with a time step of 5 µs. The membrane potential of each compartment was allowed to equilibrate to steady state before stimulation was applied. Activation was defined as membrane potential crossing 0 mV with positive slope in at least 3 compartments within the cell, initiating an action potential.

In addition to the TMS-induced E-field distributions, neural simulations were also performed using uniform E-fields and TMS waveforms, following the quasi-uniform field assumption of transcranial stimulation^67^. The extracellular potential *v*_e_ at each compartment with position (*x,y,z*) was calculated with

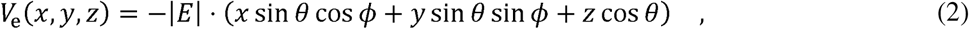

where the direction of the uniform E-field was given by polar angle *θ* and azimuthal angle *Φ*, in spherical coordinates with respect to the somatodendritic axes (Figure 1A–B), and the potential of the origin (soma) was set to zero^23^. We applied the MagProX100 monophasic TMS pulse waveform with a uniform E-field distribution at directions spanning the polar and azimuthal directions in steps of 15° and 10°, respectively, for a total of 398 directions, and we converted the spherical distribution of thresholds into 2D threshold–direction maps using the Mollweide projection (Figure 1C–D). The overall sensitivity of each model neuron to E-field direction was quantified by the *threshold anisotropy*, defined as the ratio of the maximum and minimum thresholds across all E-field directions. Additionally, the sensitivity to specific E-field directions for each model neuron was quantified by taking the average threshold for E-fields directed transverse (60° < *θ ≤* 120°) and approximately parallel to the somato-dendritic axis—either outward, towards the pial surface (0 ≤ *θ* ≤ 60°), or inward, towards the white matter (120° < *θ* ≤ 180°)—after averaging thresholds across all azimuthal rotations within the relevant polar angles.

### Neural activation thresholds and time constants

For the coupled FEM E-field and cortical layer model, single neuron thresholds were determined by scaling the coil current’s rate of change at pulse onset—which is proportional to the stimulator voltage—using a binary search algorithm to determine the minimum intensity necessary to elicit an action potential with 0.05 A/µs accuracy. For the uniform E-field simulations, threshold E-field magnitudes were obtained with 0.05 V/m accuracy.

Experimentally measured motor threshold (MT) quantifies the stimulus intensity that activates a population of corticospinal neurons, indirectly for most TMS coil configurations, with sufficient excitatory input to drive motoneurons innervating the muscle of interest. While our model cannot predict corticospinal activity or muscle excitation, we devised an approximate representation of MT for the first dorsal interosseous (FDI) muscle. We identified a putative region corresponding to the FDI representation that included both the gyral lip and upper sulcal wall (Figure 2A, Figure 6), where corticomotoneuronal projections to the corresponding spinal motoneurons originate^35^ and finger-tapping evokes functional MRI activation^7^. Within this FDI representation, we approximated MT using the threshold to activate a given percentile of model neurons, across all positions, clones, and azimuthal rotations. We then used this threshold to estimate the neural time constant using the 30, 60, and 120 µs cTMS1 pulses and the same parameter estimation method as used in previous studies^11^. The number of cortical neurons activated at MT is unknown, so we computed time constants using FDI population thresholds for a range of percentiles: 2.5, 5, 10, 25, and 50%.

### Code and data availability

The code and relevant data of this study are available from the corresponding author upon request and will be released through GitHub (https://doi.org/10.5281/zenodo.2488572).

## Acknowledgements

Preliminary results from this work were presented at the 42^nd^ Neural Interfaces Conference 2016 (Baltimore, MD, USA, Jun. 2016), 6^th^ International Conference on Transcranial Brain Stimulation (Göttingen, Germany, Sep., 2016)^68^, Society for Neuroscience’s 47^th^ Annual Meeting (Washington DC, USA, Nov., 2017), 40^th^ Annual International Conference of the IEEE Engineering in Medicine and Biology Society (Honolulu, HI, USA, Jul., 2018), and 2018 NYC Neuromodulation Conference & NANS Summer Series (New York City, NY, USA, Aug. 2018). Research reported in this publication was supported by the National Institute of Mental Health of the National Institutes of Health under award numbers R01 NS088674, U01 AG050618, and 5R25GM103765. The content is solely the responsibility of the authors and does not necessarily represent the official views of the National Institutes of Health. This work was also supported by the National Science Foundation (DGF 1106401) and a research grant by Tal Medical/Neurex Therapeutics. We thank Dr. Martin Sommer for providing raw motor threshold data, and the Duke Compute Cluster team for computational support. We also thank Dr. Marc Sommer, Dr. Stefan Goetz, Dr. Luis Gomez, Dr. Andreas Neef, Dr. Lari Koponen, Guillherme Saturnino, Rena Hamdan, and Karthik Kamaravelu for their technical assistance and helpful discussions.

## Author contributions

A.V.P. and W.M.G. conceived and supervised the study. A.S.A developed the computational methods and models, performed simulations, analyzed and interpreted results, generated figures, and wrote the manuscript. B.W. contributed to data analysis and interpretation, generation of figures, and the writing and editing of the manuscript. A.V.P. and W.M.G revised the manuscript.

## Supplementary Figures

**Supplementary Figure 1.**
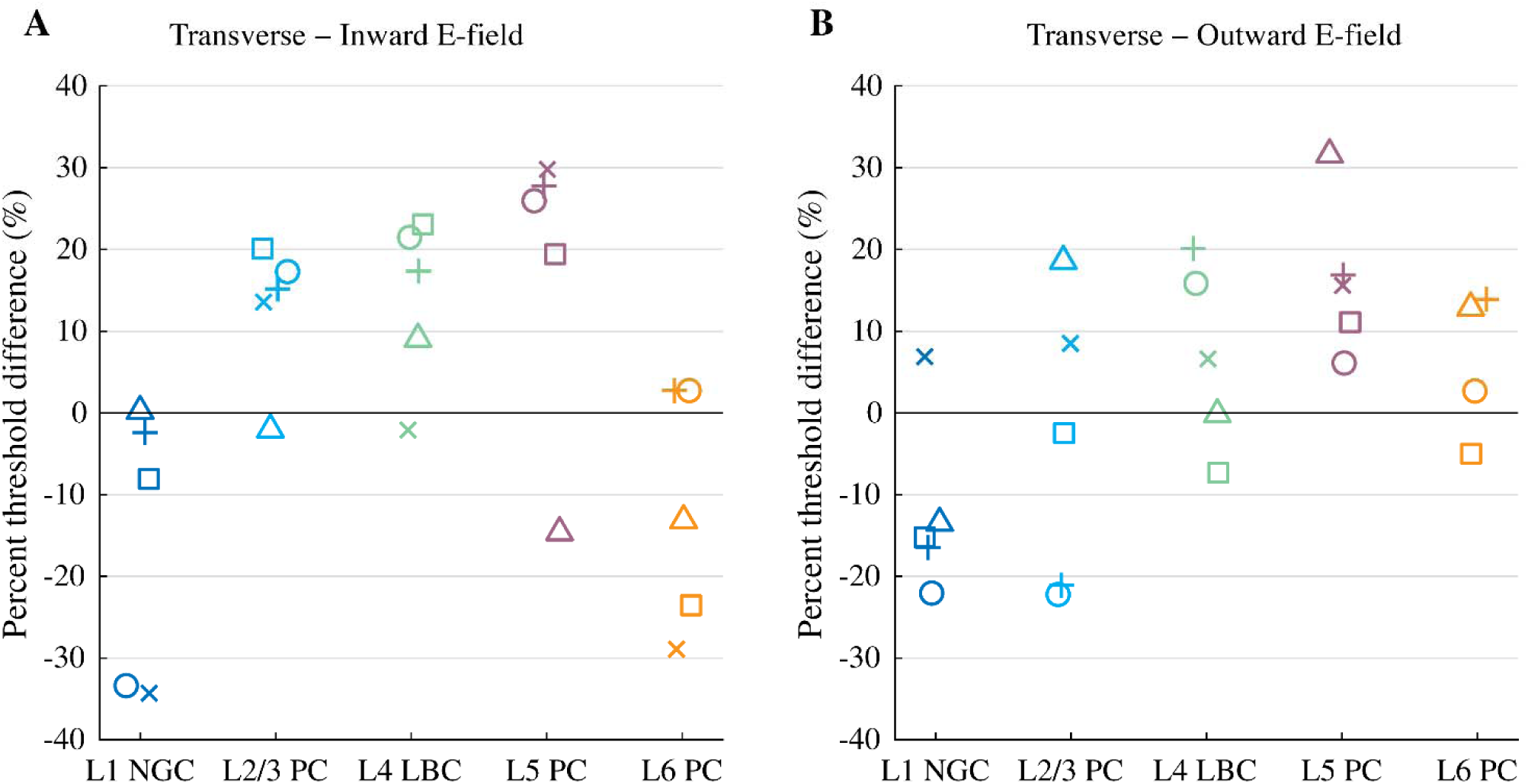
Threshold differences for transverse and inward/outward uniform E-field. Percent difference in threshold for **A)** transverse versus inward E-field and **B)** transverse versus outward E-field. Average thresholds were computed for E-fields directed transverse (60° < *θ* ≤ 120°) and approximately parallel to the somato-dendritic axis—either upwards, towards the pial surface (0 ≤ *θ* ≤ 60°), or downwards, towards the white matter (120° < *θ* ≤ 180°)—after averaging thresholds across all azimuthal rotations within the relevant polar angles. Symbols for each clone match Figure 1E.

**Supplementary Figure 2.**
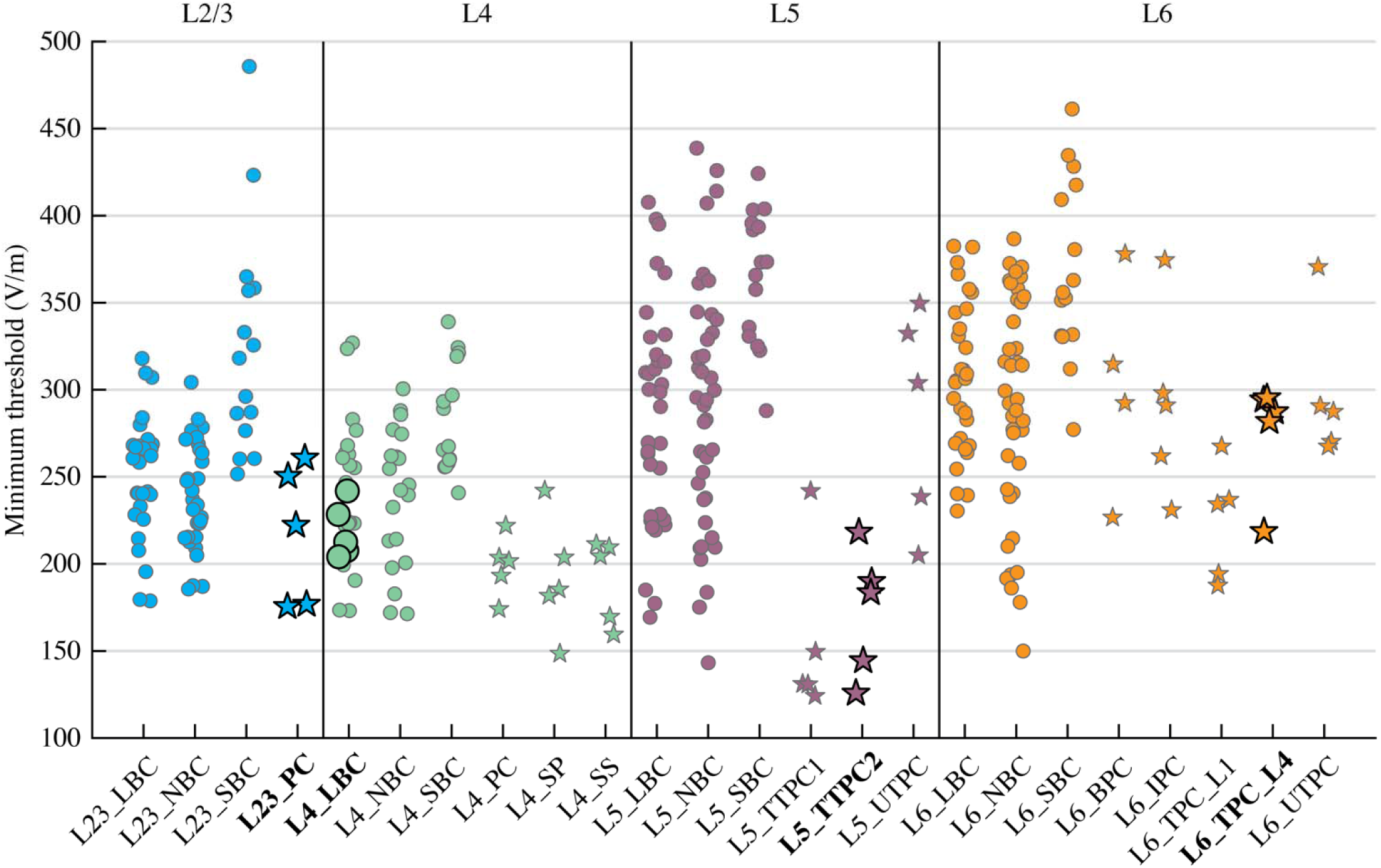
Minimum threshold for all myelinated cortical cell types in the Blue Brain library with uniform E-field. The minimum threshold was extracted from threshold–direction maps for five virtual clones of all cell types expected to possess some degree of myelination, i.e. inhibitory basket cells and excitatory cells. Inhibitory and excitatory cells are indicated by circle and star symbols, respectively. The thresholds for the model neurons that we previously published^23^ and included in the multi-scale model in this paper are shown enlarged with black outlines and bold labels. The inhibitory morphological types have more than five points because they each have multiple electrical types, with five virtual clones per morpho-electrical type. The same methods were used to modify the full library of Blue Brain library model neurons and generate threshold–direction maps, except we used polar and azimuthal steps of 30°, giving 62 total thresholds per model neuron from which the minimum was extracted, and we did not disable the main axon terminal of any of the model neurons. Additionally, some model neurons did not have a constant steady state membrane potential, so they were initialized at an approximate rest potential. The rest potential was determined for these models by simulating them for 3 sec with no stimulus and taking the mean membrane potential for the latter 2.5 sec, if the membrane potential exhibited subthreshold oscillations, or taking the final membrane potential value, if the membrane potential exhibited spontaneous firing or monotonic drift. Cell model names consist of layer (L2/3, L4, L5, or L6) and morphological-type, which are abbreviated as follows: large basket cell (LBC), nest basket cell (NBC), pyramidal cell (PC), small basket cell (SBC), star pyramidal cell (SP), spiny stellate cell (SS), small tufted pyramidal cell (STPC), thick-tufted pyramidal cell with apical trunk that bifurcated at the distal half of the apical dendrite (TTPC1) or proximal half of the apical dendrite (TTPC2), untufted pyramidal cell (UTPC), inverted pyramidal cell (IPC), bipolar pyramidal cell (BPC), and tufted pyramidal cell with apical dendrite terminating in L1 (TPC_L1) or L4 (TPC_L4). Layer 1 cells are not represented because their axons are not expected to be myelinated.

**Supplementary Figure 3.**
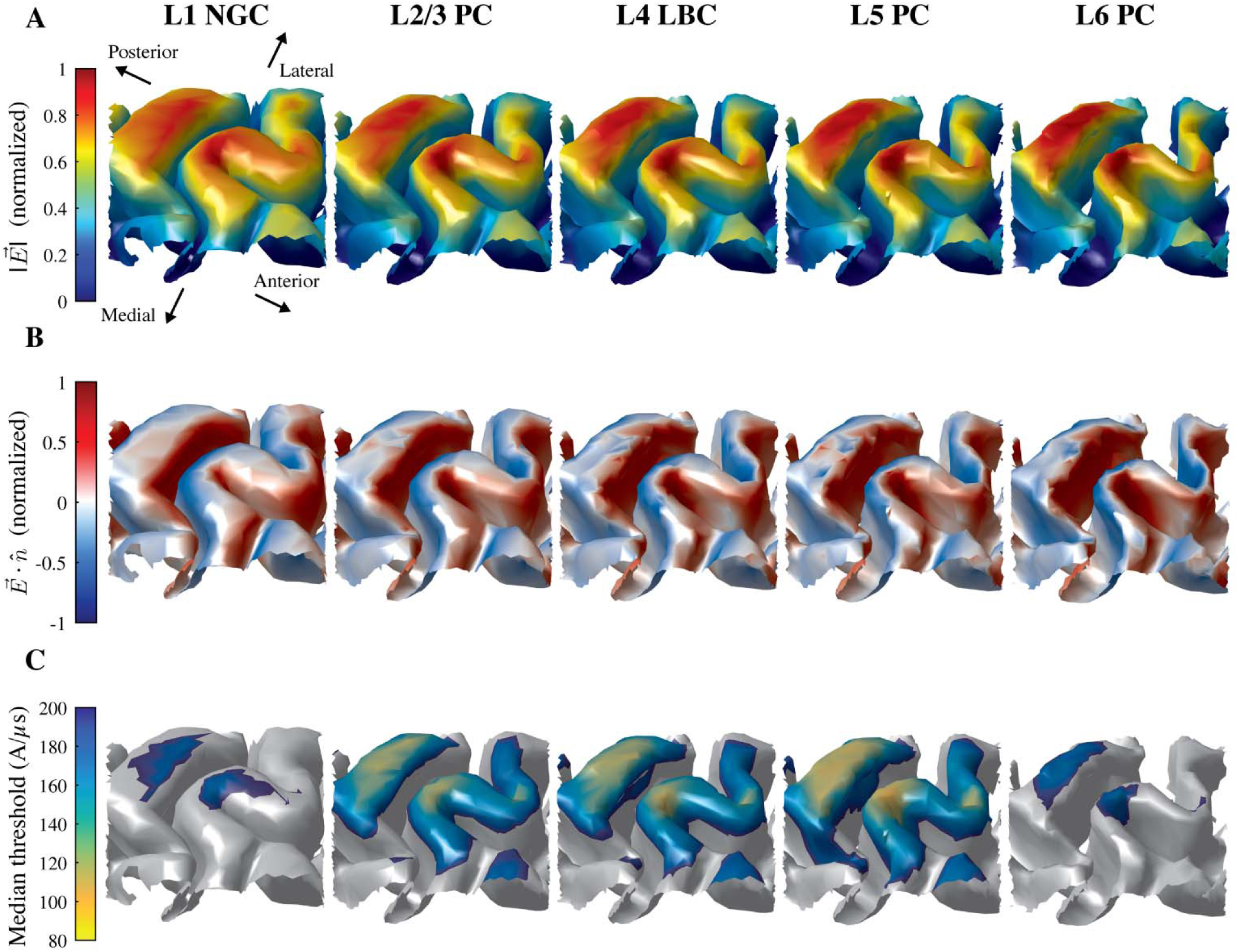
Low threshold region extends further into posterior wall of central sulcus than anterior wall. Alternative view of plots in Figure 2 from anterior side (facing posterior) of **A)** magnitude of simulated E-field (normalized to maximum layer) on layer surfaces for L1–L6, **B)** component of E-field normal to layer surfaces (normalized to maximum layer), and **C)** median thresholds (across 5 clones and 6 azimuthal rotations) for monophasic P–A simulation.

**Supplementary Figure 4.**
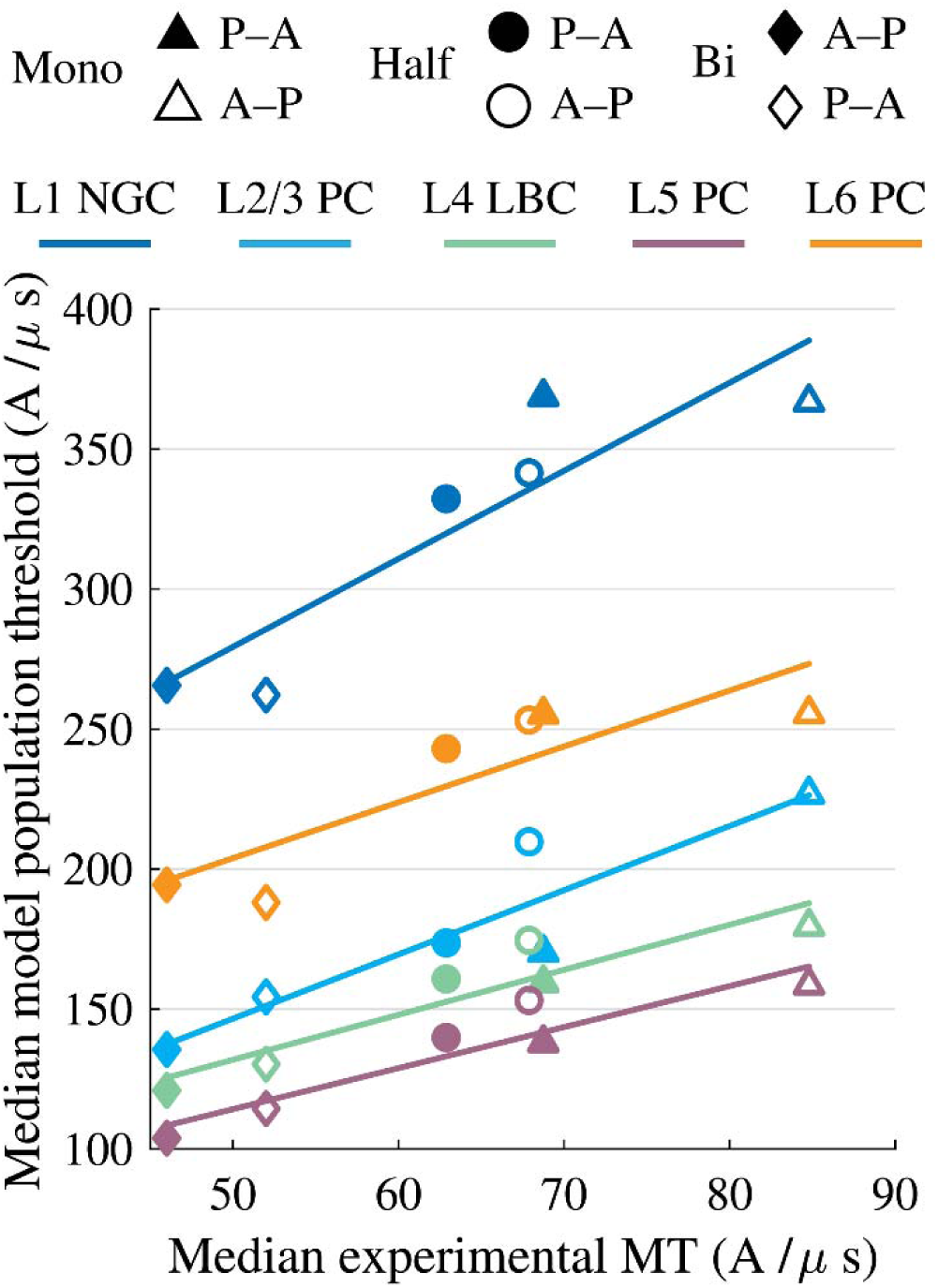
Model population thresholds correlate with experimental motor thresholds across pulse waveforms and directions. Median model population thresholds within FDI representation in each layer (same as Figure 6) plotted against median experimental motor thresholds (12 subjects)^26^. Linear regression for each layer included as solid line. *R*^2^ (*p*) values for correlations (L1–L6): 0.81 (0.015), 0.85 (0.009), 0.88 (0.006), 0.89 (0.005), 0.75 (0.025).

**Supplementary Figure 5.**
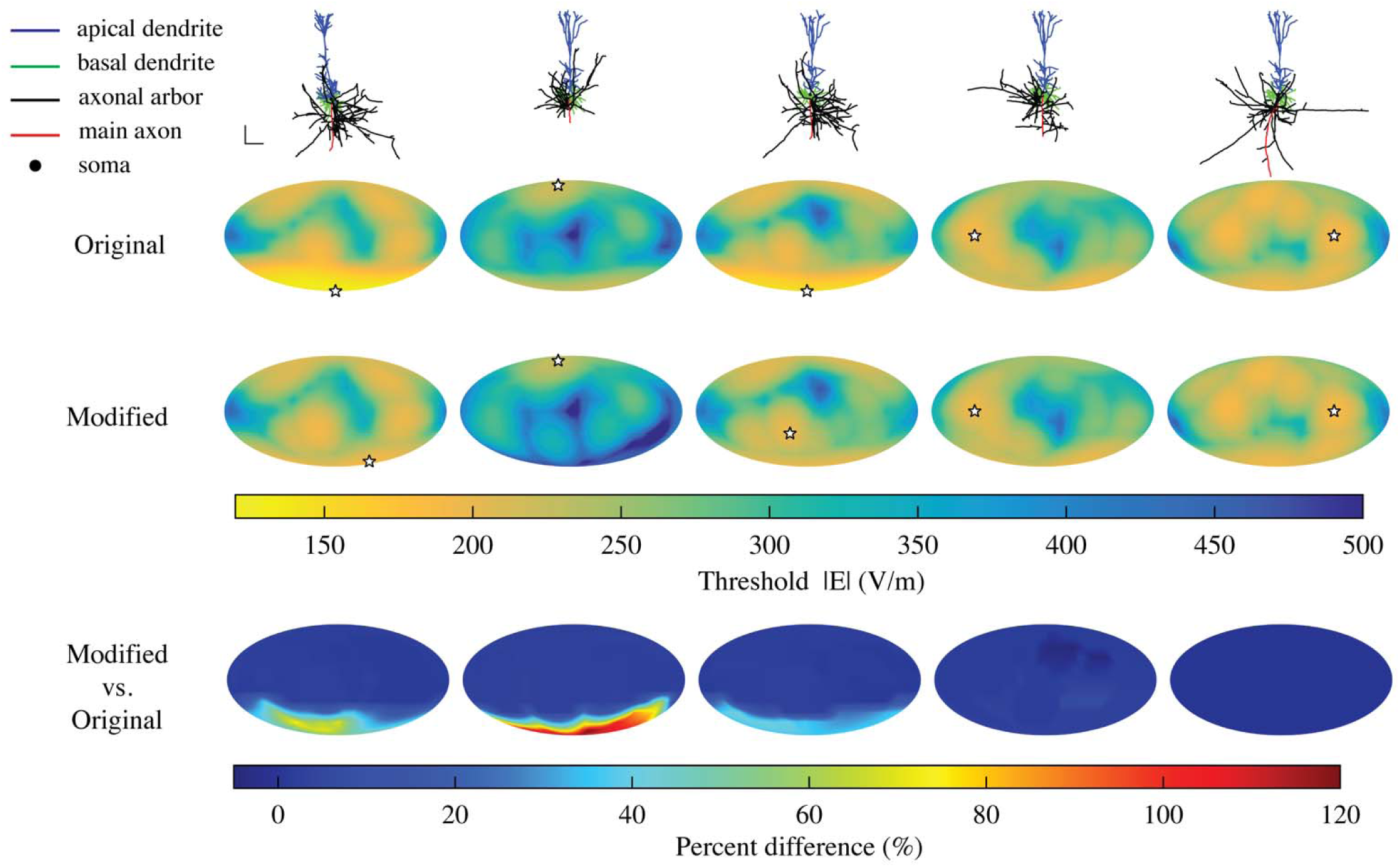
Effect of disabling main axon terminal of L5 pyramidal cells on threshold maps. Each row depicts for each L5 PC clone (from top to bottom): cell morphology; original threshold map with no modification to main axon terminal; threshold map of cell with main axon terminal disabled by setting end compartment diameter to 1000 µm; map of threshold percent differences between modified cell and original cell for each E-field direction. Notice the significant increase in thresholds for downward E-field directions in cells 1– 3 when the main axon terminal is excluded as an activation-capable site.

**Supplementary Figure 6.**
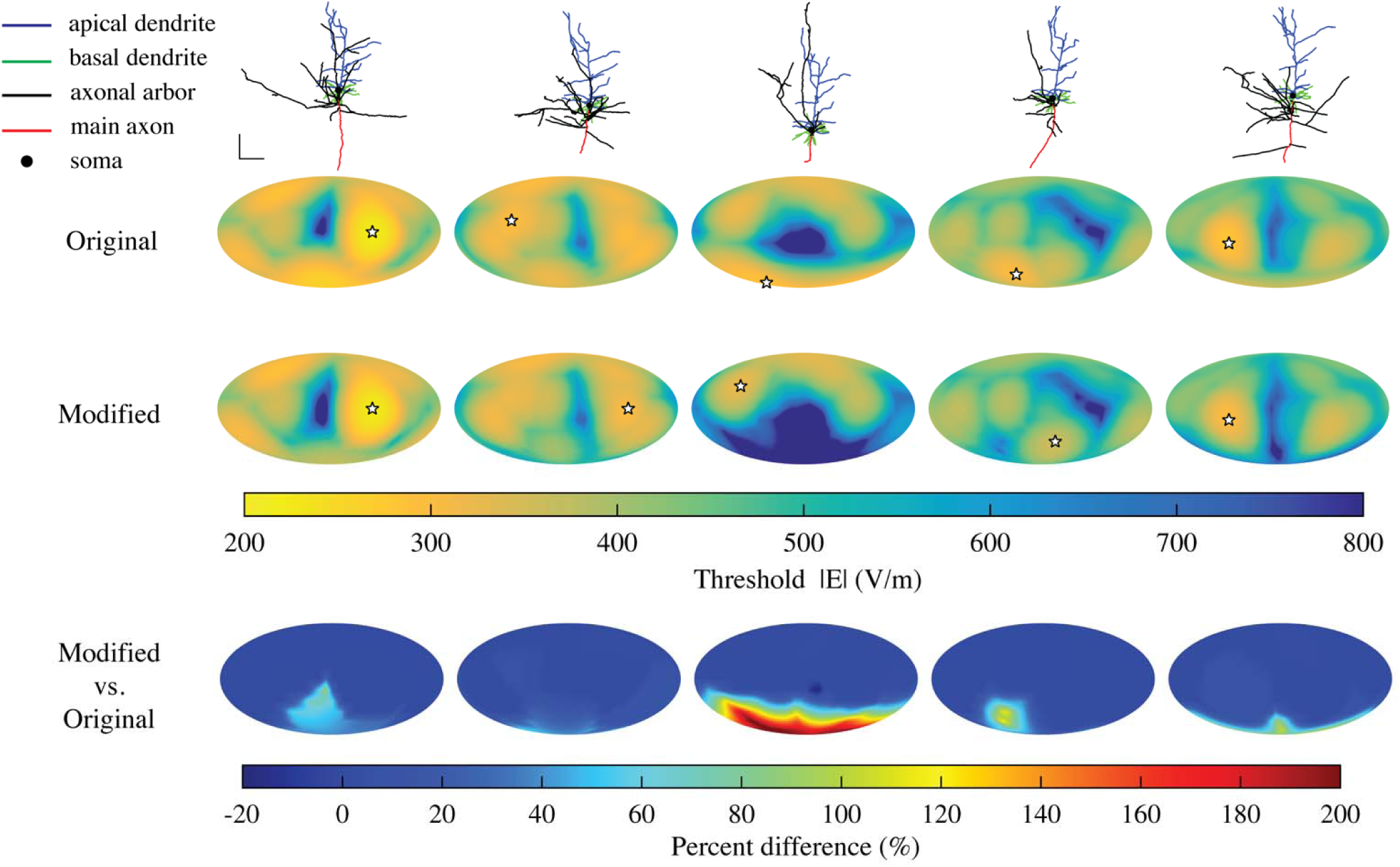
Effect of disabling main axon terminal of L6 pyramidal cells on threshold maps. Each row depicts for each L6 PC clone (from top to bottom): cell morphology; original threshold map with no modification to main axon terminal; threshold map of cell with main axon terminal disabled by setting end compartment diameter to 1000 µm; map of threshold percent differences between modified cell and original cell for each E-field direction.

